# The effects of chronic neuropathic pain states on MOR-agonist induced antihyperalgesic-like effects and rate suppressant effects

**DOI:** 10.1101/2025.08.05.668770

**Authors:** Gwendolyn E. Burgess, Hailey Y. Rizk, Shane M. Sanghvi, Emily M. Jutkiewicz

## Abstract

Chronic neuropathic pain affects 6.9-10% of the U.S. population. While mu opioid receptor (MOR) agonists are not first-line treatments for chronic pain, previous data suggests that ∼70% of chronic neuropathic pain patients receive MOR agonist treatment. Previous studies demonstrated decreased MOR expression and activity in pain states compared with non-injured or sham controls; however, we know little about how chronic pain states may alter the antinociceptive- and antihyperalgesic-like effects of MOR agonists *in vivo*. Therefore, the goal of this study was to determine if there were significant differences in MOR agonist-induced antinociceptive- or antihyperalgesic-like effects between SNI and sham groups. MOR agonist-induced antinociceptive- and antihyperalgesic-like effects were evaluated in SNI and sham groups repeatedly over 6 months. MOR agonists induced dose-dependent antinociceptive- and antihyperalgesic-like effects in male and female rats in the expected rank order of potency (fentanyl>morphine≥nalbuphine). Over time, there were small rightward shifts in fentanyl- and morphine-induced effects; however, these rightward shifts were observed in sham and SNI groups, suggesting this occurred independent of pain state. Interestingly, in sham and SNI groups, nalbuphine-induced effects were more potent 6 months post-operatively than 3 months, suggesting the potency of nalbuphine changed over time, independent of pain state. Other, non-MOR agonist analgesics were evaluated. Collectively, these data indicate that SNI failed to alter MOR agonist-induced antinociceptive- and antihyperalgesic-like effects. Future studies should evaluate if the SNI-independent increase in sensitivity to nalbuphine is related to the partial KOR agonist activity and if SNI-induced hypersensitivity alters tolerance following repeated administration of MOR agonists.

**Significance Statement:** Many chronic pain patients are treated with opioids, but we understand relatively little about how chronic pain may alter the in vivo effects of opioid analgesics. This study demonstrates that, compared to sham groups, SNI-induced hypersensitivity did not alter the potency or efficacy of opioid analgesics, opioid-induced tolerance to analgesic-like effects, or opioid-induced rate suppressant effects in either male or female rats.

## Introduction

Chronic neuropathic pain affects 6.9-10% of Americans (Scholz, et al., 2019) and is defined as pain originating from an injury or disease state affecting the somatosensory nervous system by the International Association for the Study of Pain (IASP) (Raja et al., 2019). Examples of chronic neuropathic pain include diabetic neuropathy, chemotherapeutic induced neuropathy, or post-stroke pain (Scholz et al., 2019; Rice, Smith, & Blyth, 2016; van Hecke et al., 2014).

First-line treatments for neuropathic pain include antidepressants (e.g., tricyclic antidepressants or serotonin and norepinephrine reuptake inhibitors) and anticonvulsants (pregabalin, gabapentin) (O’Connor et al., 2009; for recent review, see Bernetti et al., 2020). Many neuropathic pain patients report that these first-line treatments do not produce satisfactory relief from symptoms (Dworkin et al., 2007; Scholz et al., 2019). Therefore, second- and third-line treatments for neuropathic pain are used frequently and include topical lidocaine patches and mu opioid receptor (MOR) agonists (Cavilla et al., 2019; O’Connor et al., 2009). While there has been some debate surrounding the effectiveness of MOR agonists in treating neuropathic pain, numerous studies report MOR agonists produce analgesia in neuropathic pain states across many species, including in humans (Joubert et al., 2020). Further, some studies demonstrated MOR agonist treatment resulted in higher levels of patient satisfaction than other treatments (Widerstrom-Noga & Turk, 2003). Therefore, it is perhaps not surprising that MOR agonists are used in many chronic neuropathic pain patients with approximately 69% of patients receiving MOR agonists at some point and ∼20% of patients maintained on MOR agonists chronically (Hoffman et al., 2017). Treating chronic pain requires prolonged periods of repeated dosing with MOR agonists, so there is always some concern about development of tolerance to analgesic effects or misuse (O’Connor et al., 2009). It is well documented that repeated use of MOR agonists can produce tolerance to MOR agonist induced effects, requiring dose escalation to produce the same effects (Ballantyne and LaForge, 2007; Walker & Young, 2001). Using large doses of MOR agonists is linked to increased risk of developing an opioid use disorder (OUD) as well as risk of overdose (Ballantyne &LaForge, 2007; Hayes et al., 2020).

Although MOR agonists are frequently used in chronic neuropathic pain treatment, we understand relatively little about how chronic pain may alter the effects of MOR agonists. For example, MOR expression levels and MOR-induced G protein activation are decreased in chronic pain states (Back et al., 2006; Campos-Jurado et al, 2019; Dong et al., 2019; Hou et al., 2017; Ji et al., 1995; Kaneuchi et al., 2019; Pol et al., 2006; Thompson et al., 2018; Yamamoto et al., 2008; Zhang et al., 1998). Additionally, studies demonstrate decoupling of the MOR from the G protein in brain areas involved in reward and pain perception in chronic pain states as well as desensitization of MORs (Campos-Jurado et al., 2019; Ozaki et al., 2002). Together, these data suggest that in chronic pain states, the potency or efficacy of MOR agonist induced effects could be decreased *in vivo*.

Therefore, the goal of this study was to evaluate the potency and effectiveness of MOR agonist-induced antihyperalgesic-like effects and rate decreasing effects before induction of chronic neuropathic pain as well as over the course of 6 months of chronic neuropathic pain. Co-examination of suppression of rates of responding and analgesic effects is useful, as highly sedating drugs can produce a false positive in pain elicited tasks such as mechanical hypersensitivity (Stevenson et al., 2006; Negus et al., 2006). For these studies, we used the spared nerve injury (SNI) model (or sham control) because it produces a persistent hyperalgesic-like state that persists for at least 8 months (Decosterd & Wolfe, 2000; Erichsen & Blackburn-Munro, 2008). In addition, we evaluated these effects in in both male and female rats since few studies have evaluated the effects of MOR agonists in the SNI model though women are more likely to experience neuropathic pain than men (Bouhassira, et al., 2008).

## Methods

### Animals

All rats were at least 8 weeks old at the start of the experiment. Rats were purchased from Envigo (Indianapolis, IN), and after delivery, were given one week to habituate to the animal housing facility. They were housed in standard cages with corncob bedding with water available ad libitum. There were 6 rats per condition in all experiments except one female rat was removed from the operant assay around the 4-6 month timepoint due to a prolapsed uterus. Rats were given enviropacks and wooden gnawing blocks for enrichment. The animal housing facility operates on a 12 hr light dark cycle, and all experiments were conducted during the light cycle. Rats in the mechanical hypersensitivity experiments were group-housed for the duration of the study with food and water available ad libitum. For the operant experiments, rats were single-housed and food restricted with food restriction starting 24 hr prior to the start of operant training. Female rats were maintained on ∼13 grams of standard chow, and male rats were maintained on ∼18 grams of standard chow for the duration of the study, though water was available ad libitum.

### SNI or sham surgery

Spared nerve injury surgery was performed as described by Decosterd & Woolfe (2000). Briefly, rats were deeply anesthetized with ketamine (90 mg/kg, intraperitoneal (i.p.)) and xylazine (10 mg/kg, i.p.). 5 mg/kg carprofen (subcutaneously (s.c.)) was given pre-emptively as well as 24- and 48-hr after surgery for post-operative pain relief. An incision was made in the left femoral muscle, exposing the sciatic nerve. A 2 mm section was removed from the peroneal and tibial nerves, while the sural was not manipulated. The remaining portions of the tibial and peroneal nerves were sutured to the muscle to minimize any incidence of nerve regrowth. The sham surgery consisted of only the incision into the femoral muscle without nerve manipulation. In both surgeries, the muscle and skin were closed separately with absorbable suture (5-0) and surgical staples, respectively.

### Randall Selitto

Prior to any evaluation of mechanical threshold, all rats were habituated to the hammock-like restraint over three days for 5, 10, and 15 min respectively. For mechanical threshold evaluations, the paw pressure applicator (Randall Selitto, IITC Life Sciences, Woodland Hills, CA) was applied continuously with increasing pressure to the contralateral and ipsilateral hind paws until withdrawal or retraction of the hindpaw occurred. The Analgesiometer recorded the pressure at which hind paw withdrawal or retraction from the applicator occurred. The maximum cutoff used was 300 g of pressure.

### Hot plate

Rats were placed on a 52°C hotplate (Columbus Instruments, Columbus, OH) and the latency to lick a hind paw was recorded. The maximum allowed time on the hotplate was 45 seconds, and if the rat did not lick a hind paw or jump off the hotplate by 45 seconds, the rat was removed and the latency of 45 seconds was recorded.

### Schedule Maintained Responding

#### Equipment

All experiments were conducted in 18 standard MedPC operant chambers. These chambers were equipped with two nosepokes (ENV-114BM), a pellet dispenser (ENV-203-45), and a food hopper (ENV-200R2M). The food hopper was located in the middle of the right side of the chamber with nose pokes on either side. The active nosepoke was illuminated and responding on the active nosepoke resulted in sucrose pellet delivery and illumination of the house light. The inactive nosepoke was not illuminated, and responding on the inactive nosepoke was recorded but had no scheduled consequences.

#### Operant Training

Animals were trained to respond for 45 mg unflavored sucrose pellets (Bioserv, Flemmington, NJ) on a fixed ratio (FR)1 schedule of reinforcement in 20-min training sessions. Following each reinforcer delivery, there was a 10 sec timeout. The work requirement was gradually increased to a FR5 schedule of reinforcement. Then, the pre-session blackout was gradually increased from 0-15 min, and response periods decreased from 20 to 3 min. The final procedure consisted of three, consecutive components in which each component consisted of a 15-min blackout and a 3-min response period. Responding on the active nosepoke only resulted in sucrose pellet delivery during the 3-min response period.

#### Testing

Testing was initiated after responding in all 3 components was stable across 5 consecutive days, defined as <30% in response rate between any two consecutive days. After testing was initiated at least two consecutive days of stable responding between tests were necessary. However, if responding deviated more than 30% between days, then three subsequent, consecutive days of stable responding were needed to proceed to the next test day. On average, 1-2 tests occurred per week.

### Drugs

Fentanyl, nalbuphine, cocaine and THC were purchased from Sigma Aldrich (Burlington, MA). Morphine was purchased from Henry Schien (Novi, MI). SNC80 was generously gifted by Kenner C. Rice. NTX, diclofenac, quinpirole, and EKC were purchased from Tocris (Bristol, UK). Ketamine hydrochloride and xylazine were obtained from Hospira, INC (Lake Forest, IL). Carprofen (Rimadyl) was obtained from (Zoetis, Parsippany, NJ).

Fentanyl citrate, morphine sulphate, nalbuphine hydrochloride, naltrexone hydrochloride, cocaine hydrochloride, quinpirole hydrochloride, and ethylketazocine (EKC) were dissolved in sterile saline. SNC80 was dissolved in 3-5% 1 M HCl and the remainder of the solution was sterile water. Δ-9-tetrahydrocannabidiol (THC) was dissolved in a 10:10:80 mixture of ethanol, DMSO, and sterile water. Diclofenac sodium was dissolved in sterile water. Morphine was purchased as a 50 mg/mL solution and diluted in sterile saline to relevant concentrations. THC, diclofenac, cocaine, EKC, nalbuphine, and quinpirole were given intraperitonially (i.p.), and all other drugs were given subcutaneously (s.c.).

In operant, Randall Selitto, and hot plate experiments, all drugs were administered cumulatively. In Randall Selitto and hot plate experiments, drugs were administered 5-7 days apart to minimize tolerance from repeated administration. In operant assays, opioids were not tested more than once in the same week.

### Statistical Analyses

All ANOVA analyses were conducted in SPSS version 29. Datasets with multiple timepoints were analyzed with four-way repeated measures ANOVAs with sex, time, dose, and surgical status (SNI, sham) as independent variables and time and dose as repeated measures. Datasets with only one timepoint were analyzed with three-way repeated measures ANOVAs with sex, dose, and surgical status as independent variables and dose as a repeated measure. For all ANOVAs, corrected models were used in line with Mauchly’s W criteria. Partial eta squared (η^2^_p_) was calculated as an estimate of effect size. For main effects of dose, post-hoc one-way ANOVAs were used to determine which doses were statistically different from vehicle, and these analyses were split by other significant main effects (e.g., sex, time, or surgical status).

## Results

In the absence of surgical manipulation, baseline withdrawal thresholds did not change over 6 months [no main effect of time: F(5, 50)=1.96, p=0.10, η2p=0.16] and were not different between males and females [no main effect of sex: [F(1, 10)=3.9, p=0.08, η^2^_p_ =0.28]) (Figure 1A). Importantly, withdrawal thresholds in the naïve rats (Fig. 1A) were similar to those observed in the sham-treated rats (Fig. 1B).

**Figure 1.**
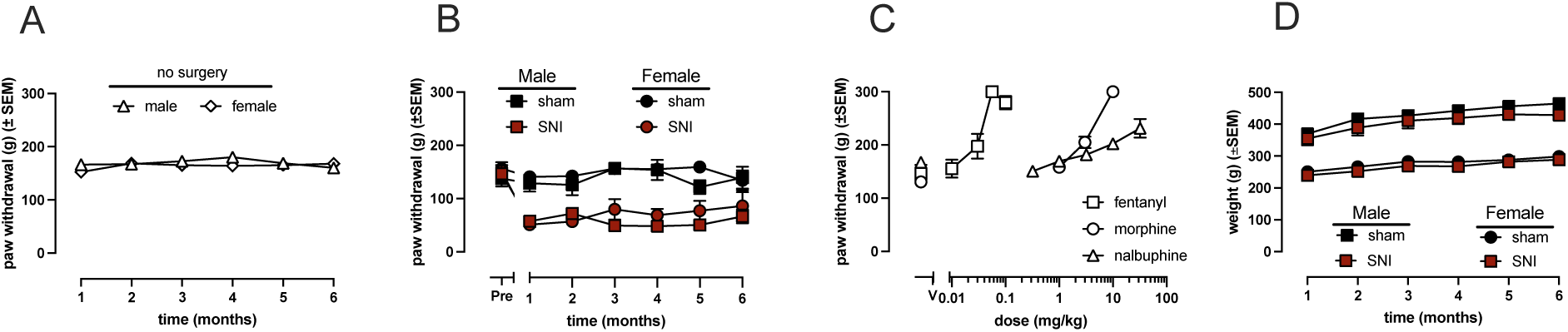
Paw withdrawal thresholds in the absence or presence of SNI-induced hypersensitivity, pre-surgical testing of opioid analgesics, and animal body weights. Paw withdrawal thresholds were unchanged in naïve male and female rats over 6 months (A). In both males and females, following sham surgery, there was no significant change in paw withdrawal thresholds compared to pre-surgical values; however, following SNI surgery, there was a significant reduction in paw withdrawal thresholds that persisted for 6 months (B). Fentanyl, morphine, and nalbuphine produced dose dependent increases in paw withdrawal thresholds in all rats prior to surgery (should probably go before B), and data are plotted with male and females collapsed (C). There is no difference in the amount of weight gained between sham and SNI animals in either males or females over 6 months (D).

As shown in Figure 1B, paw withdrawal thresholds were lower in the SNI groups than the sham groups [main effect, surgical status: F(4.02, 120)=8.09, p<0.001, η^2^_p_ =0.29] and altered over time [main effect, time: F(4.02, 120)=11.52, p<0.001, η^2^_p_ =0.37]. Specifically, there was a decrease in paw withdrawal thresholds following SNI, but not sham surgery, as evidenced by a time by surgical status interaction [F(4.02, 120)=8.09, p<0.001, η^2^_p_ =0.29] Lastly, in both sham and SNI groups, there was no sex difference in paw withdrawal thresholds over time reflected by a no main effect of sex [F(1, 20)=0.006, p=0.94, η^2^_p_ =0] and a no interaction of time and sex [F(4.02, 120)=1.48, p=0.22, η^2^_p_ =0.07] (Fig. 1B).

To further understand the changes in paw withdrawal over time in both sham and SNI groups (Fig. 1B), a one-way ANOVA was run, split by surgical status. The one-way ANOVA was not significant in the sham group [F(6, 77)=1.10, p=0.37, η^2^_p_ =0.08]. Post-hoc analyses indicated that no post-surgical timepoint was significantly different than the pre-surgical value (all p’s >0.86). The one-way ANOVA in the SNI group was significant, suggesting that the paw withdrawal thresholds changed over time [F(6, 77)=17.17, p<0.001, η^2^_p_ =0.57]. All timepoints were significantly different than pre-surgical baselines (p’s<0.001).

Prior to surgery, fentanyl, morphine, and nalbuphine dose-dependently increased paw withdrawal thresholds prior to surgery (Fig. 1C). Prior to surgery, fentanyl dose dependently increased paw withdrawal thresholds, reflected by a main effect of dose [F(2.82, 56.40)=336.33, p<0.001, η^2^_p_ =0.944]. There was no main effect of sex [F(1, 20)=0.03, p=0.80, η^2^_p_ =0.003] or surgical status [F(1, 20)=0.11, p=0.75, η^2^_p_ =0.002] in fentanyl-induced antinociceptive-like effects. All interaction terms failed to reach significance (p’s>0.12). Morphine dose dependently increased paw withdrawal thresholds prior to surgery, reflected by a main effect of dose F(1.75, 35.04)=124.86, p<0.001, η^2^_p_ =0.86] (Fig. 1C). There was no main effect of sex [F(1, 20)=0.007, p=0.93, η^2^_p_ =0] or surgical status [F(1, 20)=0.006, p=0.94, η^2^_p_ =0]. All interaction terms failed to reach significance (p’s>0.64). Nalbuphine also dose-dependently increased paw withdrawal thresholds prior to surgery, reflected by a main effect of dose [F(5, 100)=80.28, p<0.001, η^2^_p_ =0.8]. There was no main effect of sex [F(1, 20)=0.06, p=0.47, η^2^_p_ =0.03] or surgical status [F(1, 20)=0.35, p=0.56, η^2^_p_ =0.02]. All interactions failed to reach significance (p’s>0.13).

As shown in Figure 1D, SNI-induced hypersensitivity did not alter weight gain in male or female rats as compared to the sham groups. There was a main effect of sex, such that females weighed less than males, as expected [F(1, 19)=146.48, p<0.001, η^2^_p_ =0.89]. There was a significant main effect of time [F(2.5, 47.48)=69.69, p<0.001, η^2^_p_ =0.79], such that rats gained weight over time. There was also a significant interaction of time and sex [F(2.5, 47.48)=6.85, p<0.001, η^2^_p_ =0.27], reflecting differences in weight gain by sex. Interestingly, there was no main effect of surgery status [F(1, 19)=0.04, p=0.85, η^2^_p_ =0.002] and no significant interaction between time and surgical status [F(2.5, 47.48)=6.85, p<0.001, η^2^_p_ =0.27], suggesting that SNI and sham groups gained weight over time to the same extent. All other interaction terms failed to reach significance (p’s>0.10).

Fentanyl dose dependently increased paw withdrawal thresholds, supported by a main effect of dose [F(2.4, 46.21)=477.88, p<0.001, η^2^_p_ =0.96] (Fig. 2A, D). There was a main effect of surgical status [F(1, 19)=205.02, p<0.001, η^2^_p_ =0.92]. Further, there was a significant dose by surgical status interaction [F(2.4, 46.21)=24.48, p<0.001, η^2^_p_ =0.56]. These data indicate that paw withdrawal thresholds were lower in SNI as compared with sham rats, as expected, and suggest that fentanyl produced a larger increase in withdrawal thresholds in SNI rats as compared with that observed in sham groups (Fig. 2A, D). To determine which doses were significantly more effective than vehicle, one-way ANOVAs with Tukey’s post hoc analyses, split by surgical status, revealed that doses of 0.032-0.1 mg/kg significant increased paw withdrawal thresholds in sham (all p<0.001) and SNI (all p<0.001) rats. There were no main effects of sex [F(1, 19)=1.48, p=0.24, η^2^_p_ =0.07] or time [F(1.4, 26.82)=1.73, p=0.20, η^2^_p_ =0.08]; however, there was a significant time by fentanyl dose interaction [F(8, 152)= 4.36, p<0.001, η^2^_p_ =0.18] and time, dose, and sex interaction [F(8, 152)=1.86, p=0.03, η^2^_p_ =0.11]. These interactions likely reflect minor, but statistically significant, decreases in the effects of 0.056 mg/kg fentanyl over time in the male sham and SNI groups and 0.032 and 0.056 mg/kg fentanyl in the female sham group. All other interactions failed to reach significance (p’s> 0.07).

**Figure 2.**
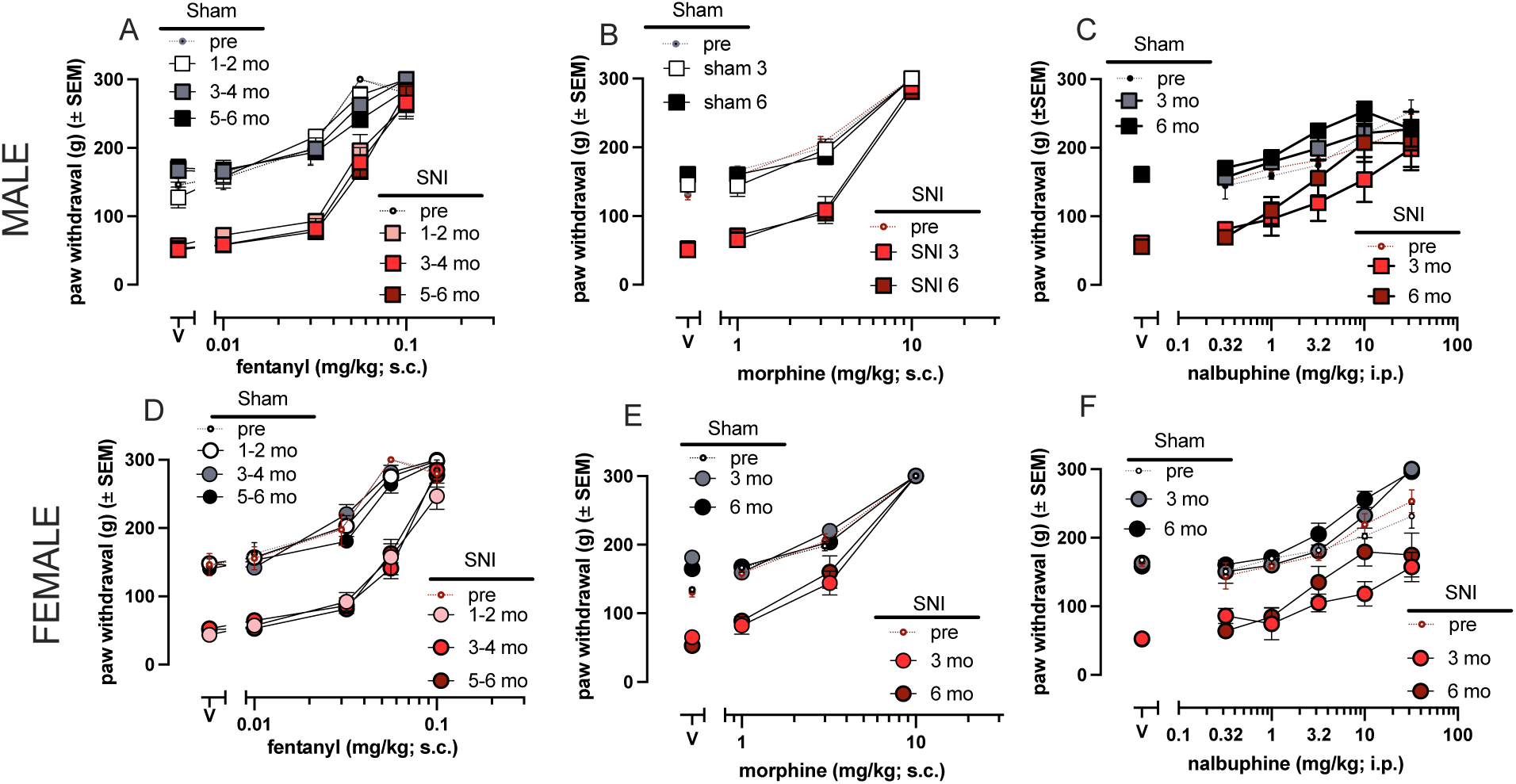
Effects of fentanyl, morphine, and nalbuphine on paw withdrawal thresholds. Fentanyl produced dose dependent increases in paw withdrawal thresholds in both males (A) and females (D) prior to surgery and following either sham or SNI; there were small, significant shifts in the dose effect curves in both sham and SNI groups, suggesting SNI did not alter fentanyl-induced effects. Morphine produced dose dependent increases in paw withdrawal thresholds in both males (B) and females (E) prior to surgery and following either sham or SNI; there were small, significant shifts in the dose effect curves in both sham and SNI groups, suggesting SNI did not alter morphine-induced effects. Nalbuphine dose dependently increased in paw withdrawal thresholds in both males (C) and females (F) prior to surgery. Following either sham or SNI, there were leftward shifts in the nalbuphine dose effect curves over time, suggesting SNI did not alter nalbuphine induced effects.

In Figure 2B and 2E, morphine produced dose dependent increases in paw withdrawal thresholds, supported by a main effect of dose [F(3, 57)=430.31, p<0.001, η^2^_p_ =0.96], with 3.2 and 10 mg/kg morphine increasing withdrawal thresholds across all treatment groups (see Table 1). There was also a significant main effect of surgical status [F(1,19)=186.74, p<0.001, η^2^_p_ =0.91] and a significant interaction between surgical status and dose [F(3, 19)=25.75, p<0.001, η^2^_p_ =0.58]. These data indicate that paw withdrawal thresholds are lower in SNI groups as compared with sham rats, as expected, and suggest that morphine produced a larger change in withdrawal thresholds in SNI rats as compared with that observed in sham groups. Morphine was slightly more potent in females, which is supported by a main effect of sex [F(1, 19)=11.99, p=0.003, η^2^_p_ =0.39]. The potency and effectiveness of morphine did not change over time, supported by no significant main effect of time [F(1, 19)=0.016, p=0.90, η^2^_p_ =0.001] or dose by time interaction [F(3, 57)=1.54, p=0.22, η^2^_p_ =0.08]. All other interactions failed to reach significance (p’s>0.18).

**Table 1.** Post-hoc testing for main effect of dose of morphine in Figure 2B, D. This table displays the output from a one-way ANOVA analyzing the main effect of dose as well as Tukey’s post-hoc testing for morphine induced antinociceptive- and antihyperalgesic-like effects in the Randall Selitto Assay (Fig. 2B, D). This one-way ANOVA was run with data collapsed across time following surgery, but data were split by sex and surgical status.

| **Table 1.** | **Main effect: dose** |  | **Post-hoc analyses** |  |  |
| --- | --- | --- | --- | --- | --- |
| **Group** | **F<sub>(3, 20)</sub>** | **p** | **P<sub>(0 vs 1)</sub>** | **P<sub>(0 vs 3.2)</sub>** | **P<sub>(0 vs 10)</sub>** |
| **Female sham** | 86.967 | **<.001** | 1 | **0.008** | **<0.001** |
| **Female SNI** | 121.998 | **<.001** | 0.62 | **0.004** | **<0.001** |
| **Male sham** | 231.163 | **<.001** | 0.56 | **<0.001** | **<0.001** |
| **Male SNI** | 101.426 | **<.001** | 0.32 | **<0.001** | **<0.001** |

As shown in figures 2C and 2F, nalbuphine dose-dependently increased paw withdrawal thresholds, which was supported by a main effect of dose [F(2.5, 45.02)=62.19, p<0.001, η^2^_p_ =0.78]. Unlike fentanyl and morphine, nalbuphine did not produce a maximal increase (e.g., at cutoff) in withdrawal threshold, consistent with a partial agonist. Interestingly, there was a main effect of surgical status [F(1, 18)=45811.32, p<0.001, η^2^_p_ =0.70] but no interaction between surgical status and dose. These analyses suggest that, as expected, paw withdrawal thresholds were lower in SNI as compared with sham rats; however, nalbuphine produced similar effects in both sham and SNI rats in contrast to that observed with fentanyl and morphine.

There was a main effect of time [F(1, 18)=25.30, p<0.001, η^2^_p_ =0.58], suggesting that nalbuphine was more effective at 6 months than 3 months (Fig. 2D, H). This is supported by a time by dose interaction [F(2.92, 52.58)=8.14, p<0.001, η^2^_p_ =0.31]. There was no main effect of sex [F(1, 18)=0.93, p=0.35, η^2^_p_ =0.05], suggesting that there were no sex differences in the effects of nalbuphine. However, there was a dose by sex by surgical status three way interaction [F(2.5, 45.02)=3.49, p=0.03, η^2^_p_ =0.16] that likely reflects the increased effect of 32 mg/kg nalbuphine in female sham group compared with the male sham group.

To determine which doses of nalbuphine were altered by time, a one-way ANOVA split by surgical status and time was run. The results are displayed in table 2; bold values are significant. 1-3 months following surgical manipulation, there was a main effect of dose and post hoc analyses determined that 10 and 32 mg/kg nalbuphine were significantly more effective than vehicle in all groups (Table 2). However, 4-6 months following surgical manipulation, 3.2-32 mg/kg nalbuphine were significantly more effective than vehicle in all groups (Table 2) suggesting that nalbuphine was somewhat more potent over time in SNI and sham groups.

**Table 2.** Post-hoc testing for main effect of dose of nalbuphine in Figure 2C, E. This table displays the output from a one-way ANOVA analyzing the main effect of dose as well as Tukey’s post-hoc testing for nalbuphine induced antinociceptive- and antihyperalgesic-like effects in the Randall Selitto Assay (Fig. 2C, E). This one-way ANOVA was run with data collapsed across sex, but data were split by time following surgery and surgical status.

| Table 2 | Main effect: dose |  |  | Post-hoc analyses |  |  |  |  |
| --- | --- | --- | --- | --- | --- | --- | --- | --- |
|  | Group | F<sub>(5, 60)</sub> | p | p<sub>(0 vs 0.32)</sub> | p<sub>(0 vs 1)</sub> | p<sub>(0 vs 3.2)</sub> | p<sub>(0 vs 10)</sub> | p<sub>(0 vs 32)</sub> |
| 1-3 mo | Sham | 18.185 | <.001 | 0.988 | 0.998 | 0.468 | <.001 | <.001 |
|  | SNI | 9.14 | <.001 | 0.78 | 0.737 | 0.084 | 0.003 | <.001 |
| 4-6 mo | Sham | 26.788 | <.001 | 0.998 | 0.688 | <.001 | <.001 | <.001 |
|  | SNI | 17.363 | <.001 | 0.989 | 0.326 | <.001 | <.001 | <.001 |

**Table 3.**
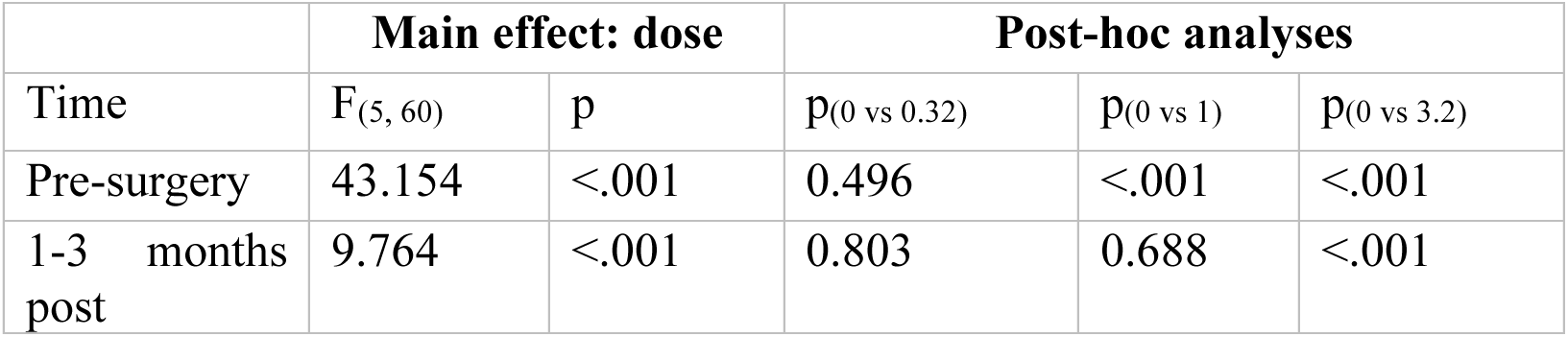
Post-hoc testing for main effect of time in Figure 7 (nalbuphine) This table displays the output from a one-way ANOVA analyzing the main effect of dose as well as Tukey’s post-hoc testing for nalbuphine-induced rate suppressant effects in the schedule maintained responding assay (Fig. 7). This one-way ANOVA was run with data collapsed across sex and surgical status to evaluate the difference in nalbuphine-induced effect prior to and after surgery.

In hotplate experiments, fentanyl dose dependently increased the latency to respond in both male and female rats (Fig. 3A, F) [main effect, dose: F(3.02, 80)=148.72, p<0.001, η^2^_p_ =0.88]. There was no main effect of surgical status [F(1, 20)=0.021, p=0.89, η^2^_p_ =0.001] or interaction between dose and surgical status [F(3.02, 80)=0.51, p=0.68, η^2^_p_ =0.14] suggesting the potency of fentanyl was not altered by surgical status. There was a significant main effect of sex [F(3.02, 80)=8.88, p=0.007, η^2^_p_ =0.31], and interaction between sex and dose [F(1, 20)=0.021, p=0.89 η^2^_p_ =0.001], indicating that fentanyl was more potent in male rats, specifically 0.032 mg/kg fentanyl.

**Figure 3.**
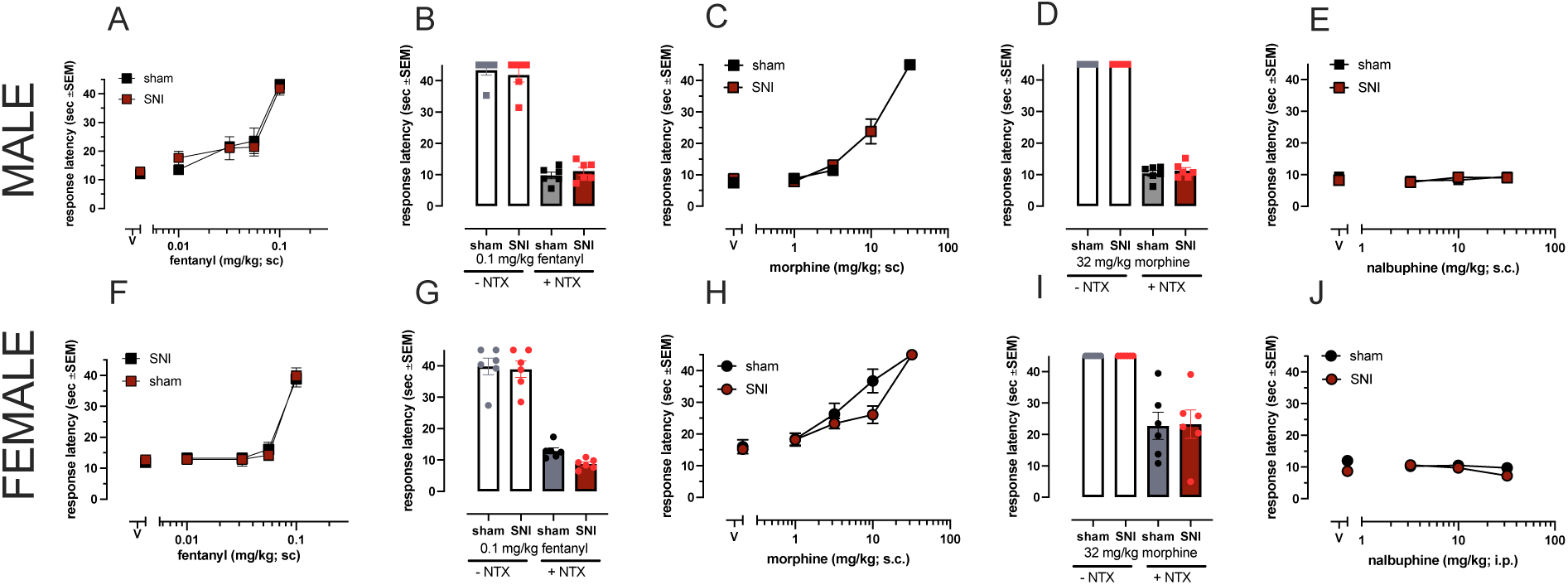
Effects of fentanyl, morphine, nalbuphine in the 52°C hot plate assay. Fentanyl produced dose-dependent antinociceptive-like effects in males (A) and females (F) regardless of surgical status. Pretreatment of naltrexone significantly reduced the antinociceptive effect of fentanyl in males (B) and females (G) regardless of surgical status. Morphine produced dose-dependent antinociceptive-like effects in males (C) and females (H) regardless of surgical status. Pretreatment of naltrexone significantly reduced the antinociceptive-like effect of morphine in males (D) and females (I), though naltrexone was significantly more effective in males. Nalbuphine failed to produce antinociceptive-like effects in either males (E) or females, regardless of surgical status (J). Data are plotted as the mean ± SEM. Data points are an average of 6 animals.

Within each sex, a one way ANOVA for dose found that the main effect of dose is significant in males [F(4, 55)=43.54, p<0.001, η^2^_p_ =0.76] and females [F(4,55)=88.03, p<0.001, η^2^_p_ =0.87]. In males, 0.032 (p=0.008), 0.056 (p=0.002), and 0.1 mg/kg (p<0.001) were significantly more effective than vehicle. In females, only 0.1 mg/kg was significantly more effective than vehicle (p<0.001). Therefore, SNI-induced hyperalgesia did not alter baselines or fentanyl-induced antinociceptive-like effects.

Morphine dose dependently increased the latency to respond in hot plate assays in both sexes (Fig. 3C, H) [main effect, dose: F(1.80, 36.02)=203.99, p<0.001, η^2^_p_ =0.91]. The main effect of dose is significant in both males [F(3, 44)=186.84, p<0.001, η^2^_p_ =0.93] and females [F(3, 44)=49.84, p<0.001, η^2^_p_ =0.77]. In males, 3.2 and 10 mg/kg morphine were more effective than vehicle (p<0.001). In females, 1 (p=0.003), 3.2 (p<0.001), and 10 (p<0.001) mg/kg morphine were more effective than vehicle. There was no main effect of surgical status [F(1, 20)=2.98, p=0.10, η^2^_p_ =0.13] and no surgical status by dose interaction [F(3, 80)=1.15, p=0.33, η^2^_p_ =0.04], further suggesting that SNI did not alter the potency of morphine. There was a significant sex difference, indicating that morphine was more potent in females than males [main effect, sex: F(1, 20)=45.56, p<0.001, η^2^_p_ =0.70]. These data were further supported by a significant interaction of sex and dose, such that 3.2 and 10 mg/kg morphine were differentially effective in males and females [F(1.80, 36.02)=7.13, p<0.003, η^2^_p_ =0.26]. All other interactions failed to reach significance (p’s> 0.17).

On the hotplate, morphine-induced antinociceptive effects were blocked by a pretreatment of 0.3 mg/kg NTX (Fig. 3D, I). There was a significant main effect of pretreatment such that NTX decreased the latency to respond in both males and females [F(1, 40)=345.13, p<0.001, η^2^_p_ =0.90]. There was also a significant main effects of sex and a significant NTX pretreatment by sex interaction, such that NTX pretreatment blocked morphine-induced antinociceptive effects to a greater extent in males (average = 10.84) than females (average = 23.34) [F(1, 40)=17.33, p<0.001, η^2^_p_ =0.30].

On the hotplate, nalbuphine did not alter response latency in either males or females up to doses of 32 mg/kg [main effect, dose [F(3, 60)=0.90, p=0.45, η^2^_p_ =0.04], regardless of surgical status [no main effect, surgical status: F(1, 20)=2.09, p=0.16, η^2^_p_ =0.09] (Fig. 3E, H). All other main effects failed to reach significance (p’s> 0.20). Further, all interaction terms failed to reach significance (p’s >0.26).

In Randall Selitto experiments, SNC80 (Fig. 4A, F) failed to alter paw withdrawal thresholds, reflected by a main effect of dose [F(3, 60)=1.86, p=0.15, η^2^_p_ =0.0]. However, there was a main effect of surgical status [F(1, 20)= 14.96, p<0.001, η^2^_p_ =0.43], suggesting that the SNI groups had lower paw withdrawal thresholds than sham groups.

**Figure 4.**
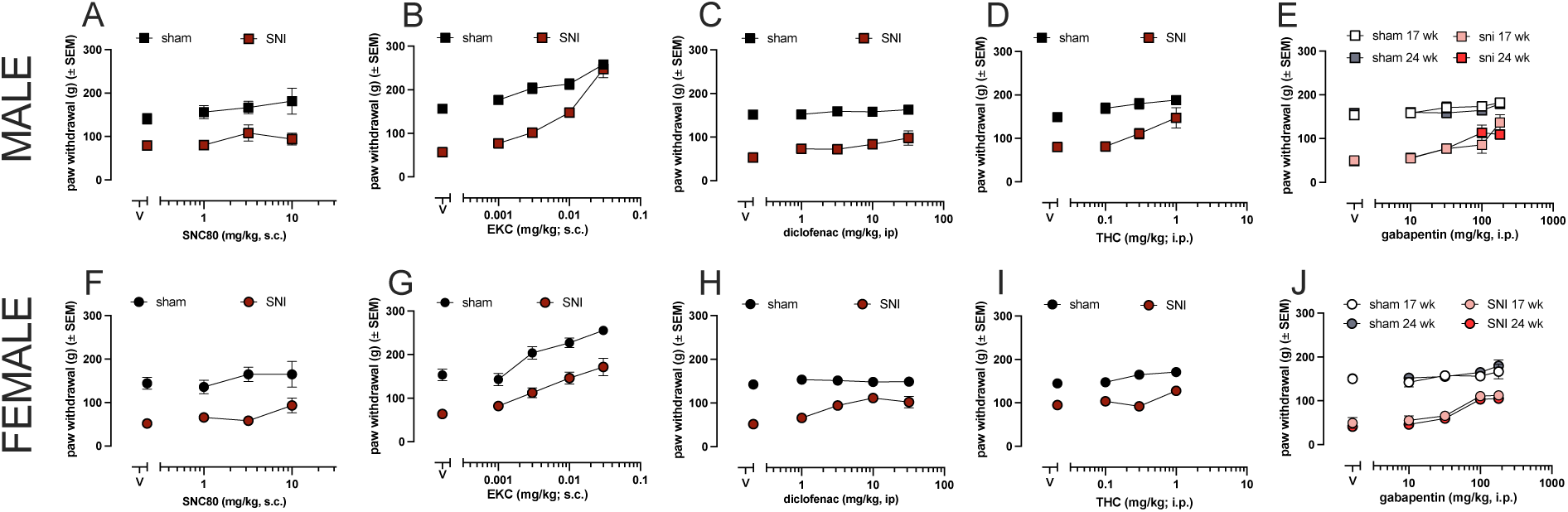
Effects of non-MOR agonist analgesics on paw withdrawal thresholds in the Randall Selitto assay. SNC80 failed to produce significant antihyperalgesic- or antinociceptive-like effects in both males (A) and females (F). EKC produced dose dependent antinociceptive- and antihyperalgesic-like effects in both males (B) and females (G). Diclofenac produced small, but significant antihyperalgesic-like effects in both males (C) and females (H) but did not produce significant antinociceptive-like effects in either sex (C, H). THC produced dose dependent antihyperalgesic-like and antinociceptive-like effects in both males (D) and females (I). Gabapentin produced dose dependent antihyperalgesic-like effects in both males (E) and females (J) but did not produce significant antinociceptive-like effects in either sex (E, J). There was no significant difference in the antinociceptive-like effects over time (E, J). Data are plotted as the mean ± SEM. Data points are an average of 6 animals.

EKC (Fig. 4B, G) dose dependently increased paw withdrawal pressures in Randall Selitto assays, reflected in a significant main effect of dose [F(2.98, 59.45)=86.57, p<0.001, η^2^_p_ =0.81]. While there was no main effect of sex, there was a significant three-way interaction between dose, surgery status, and sex [F(2.97, 59.45)=3.77, p=0.02, η^2^_p_ =0.16]. This demonstrates that EKC was approximately equally effective in the sham groups, regardless of sex, while EKC was more effective in the male SNI group than the female SNI group. Doses of 0.01 and 0.032 mg/kg EKC increased withdrawal threshold in both male and female SNI rats, though EKC was slightly more potent in sham groups with 0.0032-0.032 mg/kg producing antinociceptive-like effects in both males and females (4E, J).

In Randall Selitto experiments, diclofenac dose dependently increased paw withdrawal thresholds in SNI groups, reflected by a significant main effect of dose [F(4, 100)=7.81, p<0.001, η^2^_p_ =0.24], and there was no sex difference [main effect: sex, F(1, 20)= 0.01, p=0.92, η^2^_p_ =0.001] (Fig. 4C, H). There was a significant main effect of surgical status [F(1, 20)= 136.87, p<0.001, η^2^_p_ =0.87]. To determine which doses of diclofenac altered paw withdrawal thresholds, a one-way ANOVA was run, split by surgical status. In the sham groups, there was no main effect of dose [F(4, 55)=1.44, p=0.23, η^2^_p_ =0.095]. In the SNI groups, there was a main effect of dose [F(4, 55)=7.80, p<0.001, η^2^_p_ =0.37]. Tukey’s post hoc analysis revealed that 10 (p<0.001) and 32 (p<0.001) mg/kg diclofenac significantly increased paw withdrawal thresholds in SNI groups compared to vehicle.

Gabapentin dose dependently increased paw withdrawal thresholds in Randall Selitto assays [main effect of dose F(2.78, 55.65)=64.66, p<0.001, η^2^_p_ =0.77], and there was no sex difference [main effect of sex F(1, 20)=0.85, p=0.37, η^2^_p_ =0.04] (Fig. 4E, J). There was no change in the effects of gabapentin over time, reflected by no main effect of time [F(1, 20)=2.09, p=0.16, η^2^_p_ =0.10], no time by sex interaction [F(1, 20)=0.05, p=0.82, η^2^_p_ =0.003], and no time by surgical status interaction [F(1, 20)=2.15, p=0.16, η^2^_p_ =0.24]; however, there was a time, sex, and surgical status interaction such that gabapentin was slightly less potent in males at 3 months vs 6 months, while there were no differences in effectiveness in the female SNI group [F(1, 20)=6.36, p=0.02, η^2^_p_ =0.24]. To determine which doses of gabapentin were significantly more effective than vehicle, a one-way ANOVA was used to determine the main effect of dose in both sham and SNI groups. The one-way ANOVA was significant in sham groups [F(4,55)=2.63, p=0.04, η^2^_p_ =0.16]; however, Tukey’s post hoc analyses revealed that all doses tested failed to produce significantly different effects than vehicle. 180 mg/kg was close to the threshold for significance (p=0.06), however, the difference between the effects of 180 mg/kg and vehicle are not statistically different. In the SNI group, the one-way ANOVA found a significant main effect of dose [F(4,55)=5.03, p=0.002, η^2^_p_ =0.27], and Tukey’s post hoc analyses determined that 180 mg/kg gabapentin significantly increased paw withdrawal threshold compared to vehicle in SNI rats.

In Randall Selitto experiments, quinpirole produced dose-dependent increases in the paw withdrawal thresholds, supported by a main effect of dose [F(3.00, 59.88)=38.23, p<0.001, η^2^_p_ =0.66], with significant increases in withdrawal threshold as compared with vehicle at 0.1 mg/kg (p=0.002, sham rats) or 0.32 and 1 mg/kg (both p<0.001, SNI rats) (Fig. 5A, C). There was a main effect of surgical status as expected [F(1, 20)=47.33, p<0.001, η^2^_p_ =0.7], but there was no main effect of sex [F(1, 20)=0.06, p=0.80, η^2^_p_ =0.003]. All interaction terms failed to reach significance, including the three-way interaction (p>0.36).

**Figure 5.**
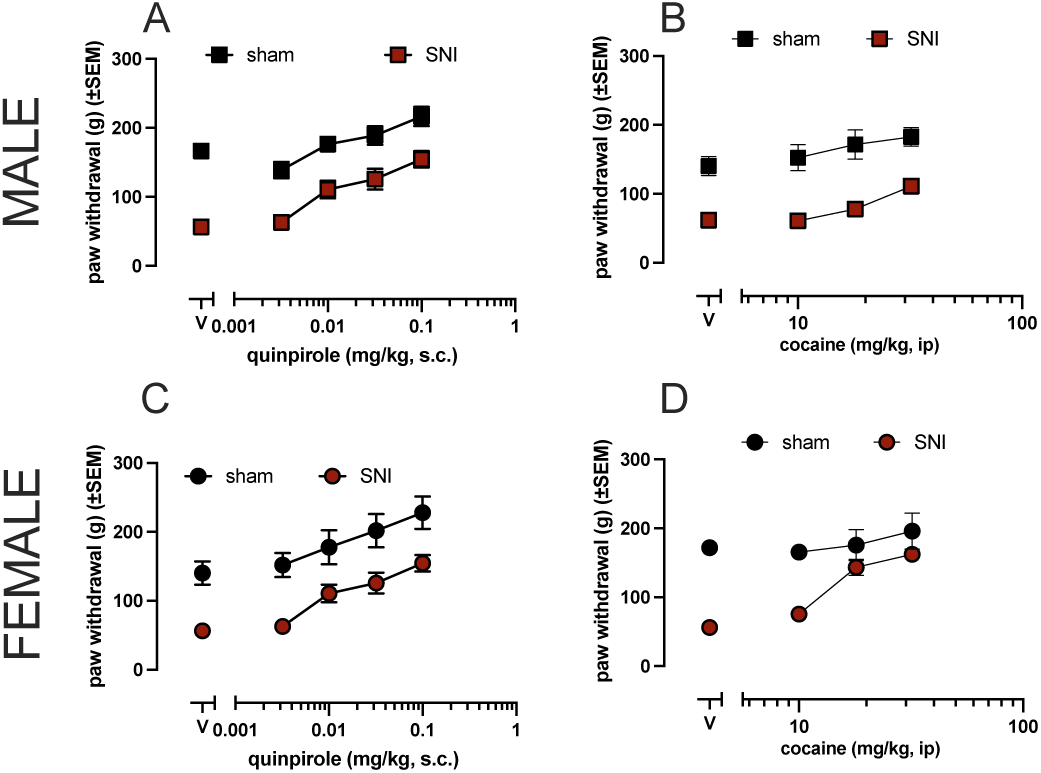
Effects of dopaminergic drugs on paw withdrawal thresholds. Quinpirole produced dose dependent antinociceptive-like and antihyperalgesic-like effects in both males (A) and females (C). Cocaine failed to produce antinociceptive-like effects in either males (B) or females (D). However, cocaine produced dose-dependent antihyperalgesic-like effects in both males (B) and females (D), though cocaine was significantly more effective in females. Data are plotted as the mean ± SEM. Data points are an average of 6 animals.

In Randall Selitto experiments, cocaine dose dependently increased paw withdrawal thresholds [main effect of dose [F(2.44, 48.84))=18.57, p<0.001, η^2^_p_ =0.48] (Fig. 5B, D). As expected, there was a significant main effect of surgical status [(F(1, 20)=72.90, p<0.001, η^2^_p_ =0.79]. There was a significant sex difference [main effect of sex [F(1, 20)=6.98, p=0.02, η^2^_p_ =0.26] such that cocaine was more effective in female rats, regardless of surgical condition. Further, there was a three way interaction between dose, surgical status, and sex [F(2.44, 48.84)=3.23, p=0.04, η^2^_p_ =0.14] such that cocaine was more effective in the female SNI group than any other group (Fig. 2D). To further analyze the effects of cocaine, one-way ANOVAs were used to compare the effects of different cocaine doses within each sex and surgical status group. There was no main effect of dose in the female sham group [F(3, 20)=0.53, p=0.67, η^2^_p_ =0.07] or male sham group [F(3, 20)=1.20, p=0.33, η^2^_p_ =0.15], suggesting that cocaine did not alter paw withdrawal thresholds in sham rats regardless of sex. There was a significant main effect of dose in the female SNI group [F(3, 20)=45.47, p<0.001, η^2^_p_ =0.87], with significant increases in withdrawal threshold at 18 and 32 mg/kg (both p<0.001). There was a significant main effect of dose in the male SNI group [F(3, 20)=8.39, p<0.001, η^2^_p_ =0.56], with significant increases in withdrawal threshold at 32 mg/kg (p=0.002).

Rates of responding were not altered by sham or SNI surgery in males (Fig. 6A) or females (Fig. 6B). As shown in figure 6, sham or SNI surgery did not alter rates of responding for sucrose pellets in male or female rats. There was no main effect of sex [F(1, 20)=0.07, p=0.79, η^2^_p_=0.004], surgery status, [F(1, 20)=2.58, p=0.12, η^2^_p_ =0.11], or time (pre, post) (Fig. 6A, B). All interaction terms failed to reach significance (p’s>0.14).

**Figure 6.**
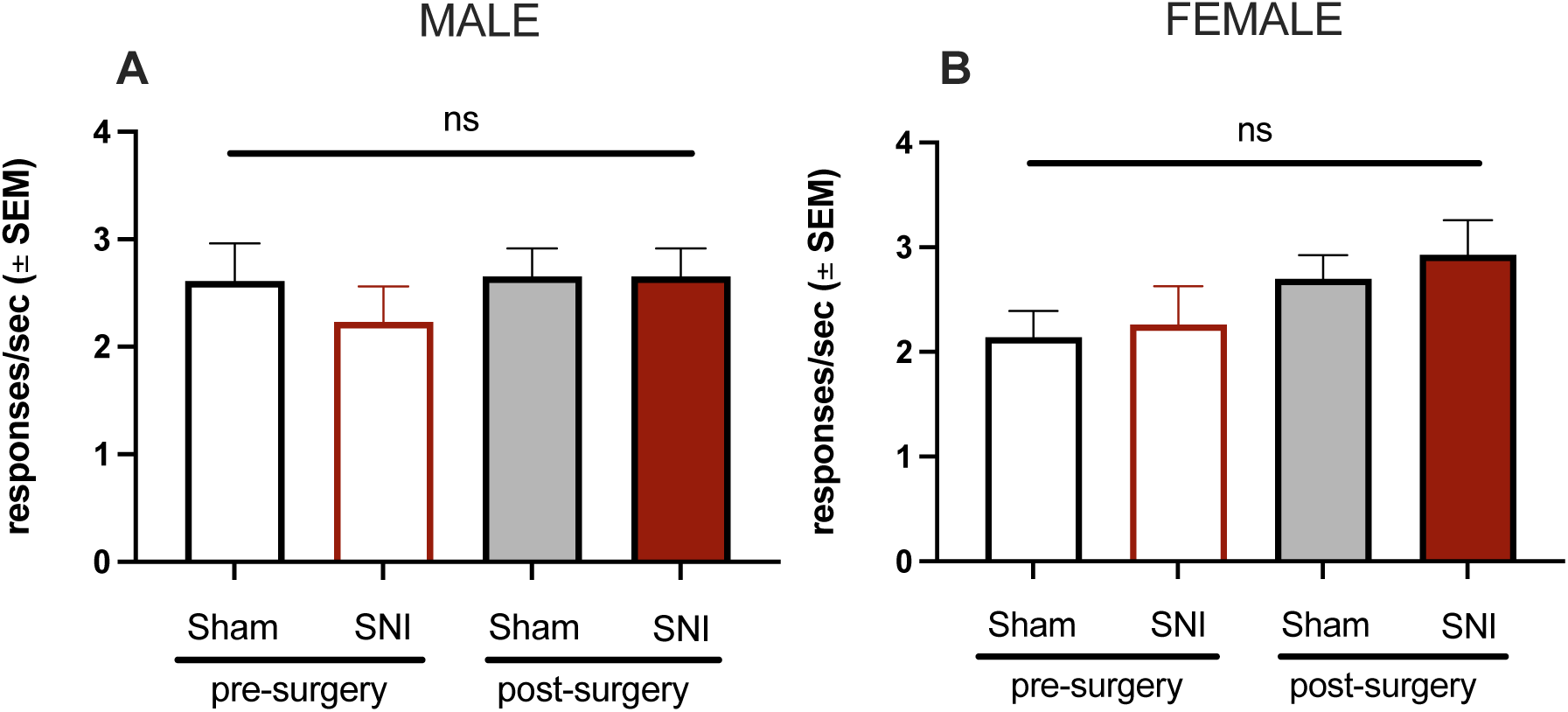
Drug free rates of responding prior to and after sham or SNI surgery. Prior to surgery, there were no differences in rates of responding between males (A) and females (B). Following sham or SNI surgery, there was no significant difference in rates of responding in either males (A) or females (B). Data are plotted as the mean ± SEM. Data points are an average of 6 animals.

The effects of fentanyl on rates of responding are shown in Figure 7. Fentanyl dose dependently decreased rates of responding in both males (Fig. 7A, B) and females (Fig. 7C, D), reflected by a significant main effect of dose [F(2.52, 37.72)=154.35, p<0.001, η^2^_p_ =0.91]. There was no main effect of time [F(2, 30)=0.87, p=0.43, η^2^_p_ =0.06], surgical status [F(1, 15)=0.002, p=0.97, η^2^_p_ =0], or sex [F(1, 15)=3.06, p=0.10, η^2^_p_ =0.17]. All interaction terms failed to reach significance (p’s >0.21). Analysis of data collapsed across time and sex show that fentanyl doses of 0.032-0.1 mg/kg significantly decreased rates of responding as compared with vehicle (Tukey’s posthoc, p<0.001). Together, these data indicate that fentanyl-induced rate decreasing effects were not altered by SNI surgery for up to six months post-injury and there were no sex differences in the rate decreasing effects of fentanyl and up to 6 months of SNI induced hyperalgesic-like states. The effects of morphine on rates of responding are shown in Figure 7. Morphine dose dependently decreased rates of responding in both males (Fig. 7A, B) and females (Fig. 7C, D). Analysis of data collapsed across time and sex show that morphine doses of 3.2 and 10 mg/kg significantly decreased rates of responding as compared with vehicle (Tukey’s posthoc, p<0.001). There was no main effect of time [F(2, 38)=2.92, p=0.07, η^2^_p_ =0.13], surgical status F(1, 19)=0.0.003, p=0.96, η^2^_p_ =0], or sex [F(1, 19)=0.28, p=0.60, η^2^_p_ =0.02]; however, there was a significant four-way interaction between time, sex, dose, and surgical status [F(6, 114)=3.14, p=0.007, η^2^_p_ =0.14]. These analyses suggest that there were likely small changes in the effectiveness of single doses of morphine across the various conditions. For example, in males, 3.2 mg/kg morphine was less rate suppressant 1-3 or 4-6 months following SNI surgery and 4-6 months post-surgery in the sham group. In contrast, the effects of morphine are relatively unchanged in female rats over time. Overall, these results suggest that the rate decreasing effects of morphine were slightly less potent over time but perhaps only in male rats.

**Figure 7.**
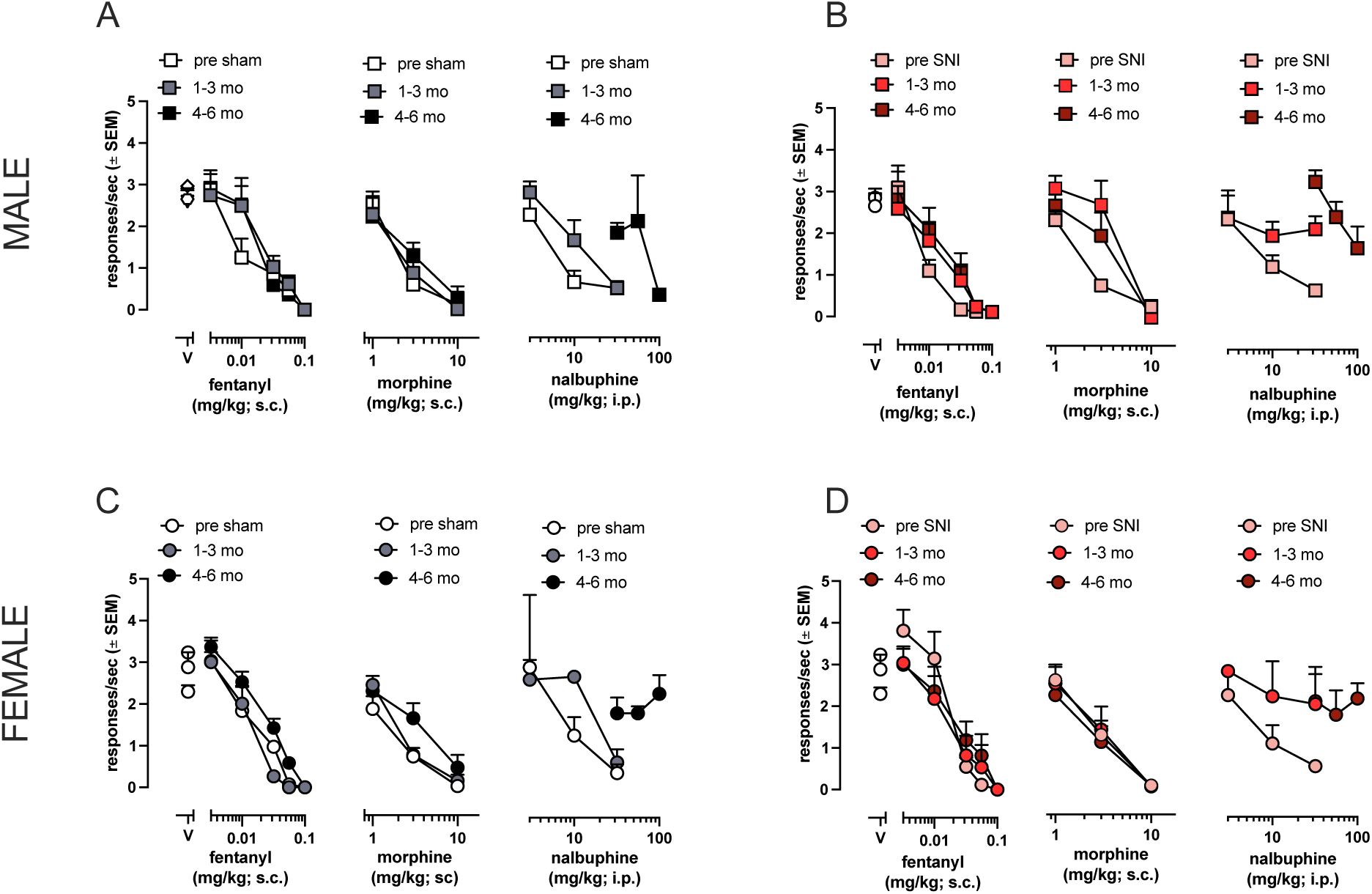
Rate suppressant effects of opioid analgesics in male and female rats prior to and 1-2 months or 3-4 months after sham or SNI surgery. Fentanyl dose-dependently decreased rates of responding prior to surgery in both males (A, B) and females (C, D). Following sham or SNI surgery, there were not significant shifts in the dose effect curves in males (A, B) or females (C, D). Morphine dose-dependently decreased rates of responding prior to surgery in both males (A, B) and females (C, D). After sham or SNI surgery, there were small, but significant shifts in the dose effect curves in both sham and SNI groups. Nalbuphine dose-dependently decreased rates of responding in males (A, B) and females (C, D). 1-3 months after surgery, nalbuphine was less potent in all groups, but still suppressed rates of responding. 4-6 months after surgery, up to 100 mg/kg nalbuphine failed to suppress rates of responding in any group except the male sham group (A). Data are plotted as the mean ± SEM. Data points are an average of 6 animals.

The effects of nalbuphine on rates of responding are shown in Figure 7. Since nalbuphine dose effect curves established 4-6 months post-surgery were substantially different from those measured pre-surgery and 1-3 months post-surgery, data analyses only directly compared nalbuphine dose effect curves established pre-surgery and 1-3 months post-surgery. Nalbuphine dose-dependently decreased rates of responding, as supported by a main effect of dose [F(2.17, 26.09)=41.15, p<0.001, η^2^_p_ =0.77]. Further there was not a significant dose by surgical status interaction [F(3, 26)=0.56, p=0.67, η^2^_p_ =0.04]. There was no main effect sex or surgical status, suggesting that there were no differences in the rate decreasing effects of nalbuphine between females and males and between sham and SNI conditions. There was a main effect of time [F(2.34, 28.05)=3.82, η^2^_p_ =0.24] and a significant interaction of time and nalbuphine dose [F(2.17, 26.09)=41.15, p<0.001, η^2^_p_ =0.77]. These data indicate that nalbuphine became less potent over time in both males and females independent of surgical status. Interestingly, 4-6 months after induction of sham (7A, C) or SNI (7B, D), nalbuphine was much less effective in decreasing rates of responding, with doses up to 100 mg/kg nalbuphine producing minimal, if any, rate decreasing effects. Overall, these data indicate that there were small changes in the rate decreasing effects of fentanyl and morphine and large rightward shifts in the effects of nalbuphine over time, and time, rather than nerve injury, produced significant changes in the rate decreasing effects of these MOR agonists. Overall, nalbuphine induced rate suppressant effects were subject to large changes over time, however, these changes are not explained by SNI-induced hypersensitivity.

As shown in Figure 8, cocaine dose-dependently decreased rates of responding (Fig. 8A-D) prior to surgery, 1-3 months, and 4-6 months after both SNI or sham surgeries supported by a main effect of dose [F(2.05, 22.59)=63.05, p<0.001, η^2^_p_ =0.85], with significant rate decreasing effects at 18 and 32 mg/kg as compared with vehicle (p<0.001 both). All other main effects and interactions failed to reach significance (p’s >0.16). This suggests that cocaine induced rate suppressant effects were not altered over time or by nerve injury.

**Figure 8.**
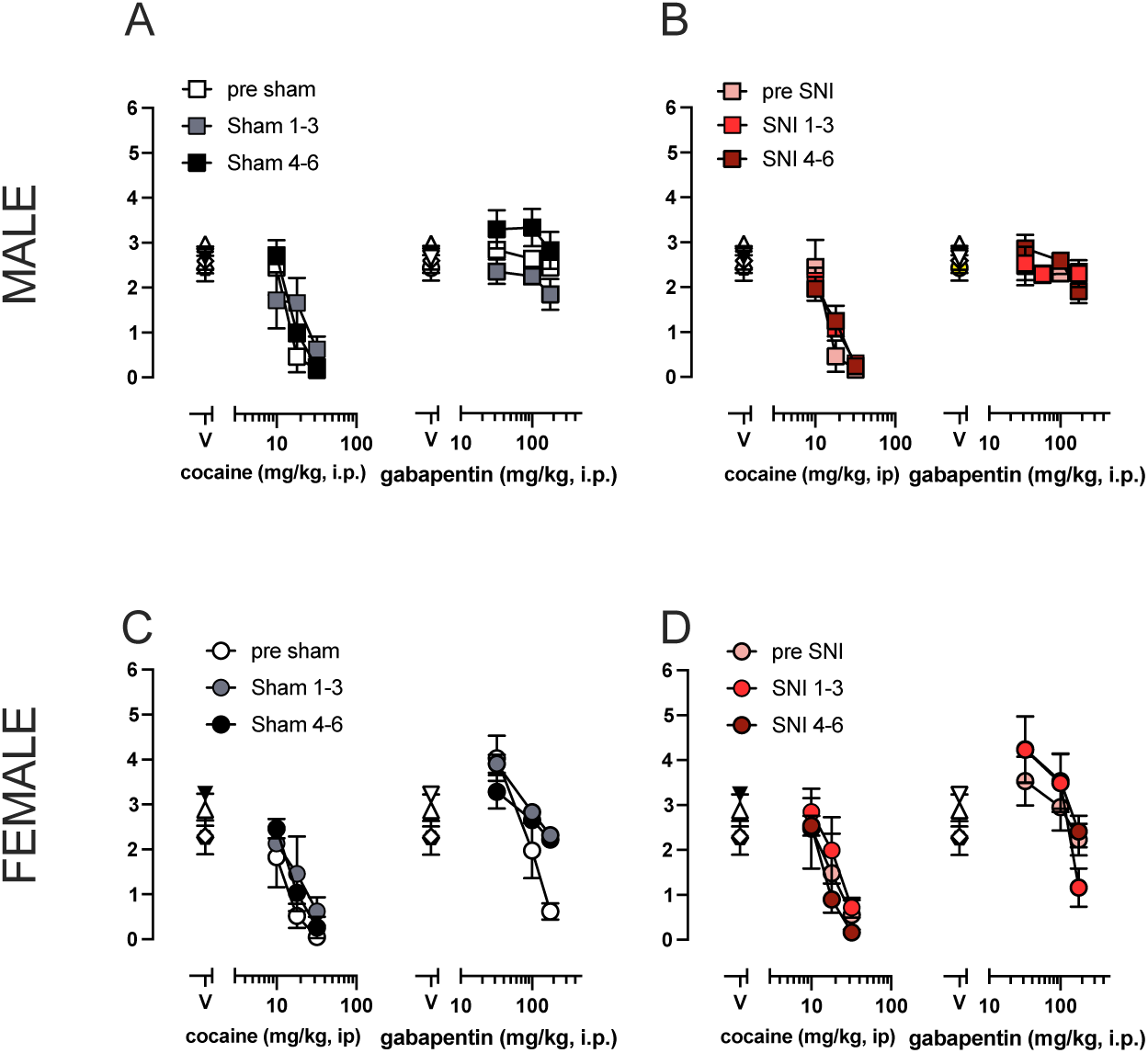
Rate suppressant effects of cocaine and gabapentin in male and female rats prior to and 1-2 months or 3-4 months following sham or SNI surgery. Prior to sham or SNI surgery, cocaine dose dependently decreased rates of responding in both males (A, B) and females (C, D). After sham or SNI surgery, there were no significant shifts in the dose effect curves in either males (A, B) or females (C, D). Gabapentin failed to alter rates of responding in male rats prior to either sham (A) or SNI surgery (B). There were no significant changes following sham (A) or SNI (B) surgery such that gabapentin did not alter rates of responding in male rats. In females, prior to and after either sham (C) or SNI (D) surgery, low doses of gabapentin (32 mg/kg) increased rates of responding compared to vehicle while larger doses (180 mg/kg) significantly decreased rates of responding compared to vehicle. There were no significant differences between sham (C) and SNI (D) groups or in dose effect curves over time following surgery. Data are plotted as the mean ± SEM. Data points are an average of 6 animals.

As shown in Figure 8(A-D), gabapentin decreased rates of responding, reflected by a main effect of dose [F(3, 26.15)=12.12, p<0.001, η^2^_p_ =0.50]. There were no main effects of time [F(2, 24)=0.41, p=0.67, η^2^_p_ =0.03] or surgical status [F(1, 12)=4.49, p=0.06, η^2^_p_ =0.27], suggesting that generally, gabapentin-induced rate decreasing effects are not altered by nerve injury or over time. However, there was a main effect of sex [F(1, 12)= 10.29, p=0.008, η^2^_p_ =0.46], a sex by time interaction [F(2, 24)=4.82, p=0.02, η^2^_p_ =0.29], and a sex by time by surgical status three-way interaction [F(6, 72)=4.03, p=0.002, η^2^_p_ =0.09], indicating that gabapentin did not decrease rates of responding in male rats up to a dose of 180 mg/kg and that the gabapentin decreased response rates to a lesser extent over time in female sham rats as compared with female SNI rats.

## Discussion

Previous studies have demonstrated that pain states, including neuropathic pain states, resulted in increased levels of endogenous opioid peptide release (Bruehl et al., 2012, 2013, 2014), decreased MOR expression (Back et al., 2006; Campos-Jurado et al, 2019; Dong et al., 2019; Hou et al., 2017; Ji et al., 1995; Kaneuchi et al., 2019; Pol et al., 2006; Thompson et al., 2018; Yamamoto et al., 2008; Zhang et al., 1998), and decreased MOR activation (Campos-Juardo et al., 2019; Ozaki et al., 2002). However, we know relatively little about if chronic pain states alter MOR agonist-induced antinociceptive- or antihyperalgesic-like effects over time. Therefore, the goal of this study was to evaluate MOR agonist-induced antinociceptive-like effects, antihyperalgesic-like effects, and rate suppressant effects in animals in the presence or absence of SNI-induced hypersensitivity over 6 months following surgery.

In the present study, SNI surgery induced a persistent decrease in paw withdrawal thresholds from mechanical stimuli as compared with the sham control condition, suggesting long-lasting mechanical hypersensitivity following SNI surgery (Fig. 1). Acutely, fentanyl, morphine, and nalbuphine induced dose-dependent antinociceptive- and antihyperalgesic-like effects in Randall Selitto assays (Fig. 1C, 3). Unlike fentanyl and morphine, nalbuphine failed to produce a maximum effect in the Randall Selitto assay (Fig. 2C, F), and nalbuphine failed to produce significant antinociceptive-like effects in the hot plate assay (Fig. 3). Both findings are consistent with partial MOR agonist activity. Additional non-MOR agonist drugs with previously reported analgesic-like effects were tested in Randall Selitto assays. SNC80, a DOR agonist, failed to alter paw withdrawal thresholds, but EKC, a KOR agonist, produced dose-dependent antinociceptive- and antihyperalgesic-like effects in both sexes (Fig. 4). THC, diclofenac, and gabapentin produced antihyperalgesic-like effects but not antinociceptive-like effects (Fig. 4). Quinpirole produced dose dependent antinociceptive- and antihyperalgesic-like effects, and cocaine produced antihyperalgesic-like effects (Fig. 5).

To determine if chronic neuropathic pain altered the potency and or/efficacy of MOR agonist-induced antinociceptive- or antihyperalgesic-like effects, fentanyl-, morphine-, and nalbuphine-induced effects were repeatedly assessed over 6 months following sham and SNI surgery. Over time, small shifts in the effects of individual doses of fentanyl and morphine were observed; however, this was observed *independent* of surgical status, suggesting that SNI-induced hypersensitivity does not explain the small changes in MOR agonist-induced effects and that other factors likely underly the changes in dose response curves over time in both groups.

For instance, MOR agonists can produce robust behavioral disruption, and one measure of this is decreased rates of responding. Fentanyl and morphine dose-dependently decreased rates of responding prior to surgery in all groups (Fig. 7). 1-3 months or 4-6 months after surgery, there were changes in the effectiveness of individual doses of fentanyl and morphine independent of surgical status; however, there were no large shifts in dose effect curves. These data collectively suggest that the behavioral effects of higher efficacy MOR agonists were not altered by the presence of chronic neuropathic pain states. Though previous studies have demonstrated a decrease in MOR expression or function in pain states, it is possible this decrease is not sufficient to alter the effects of higher efficacy MOR agonists.

Unlike fentanyl and morphine, nalbuphine-induced antinociceptive- and antihyperalgesic-like effects were more potent 6 months following surgery than 3 months in both male and female rats (Fig. 2C, F). Prior to surgery, nalbuphine dose-dependently decreased rates of responding in all groups (Fig. 2). 1-3 months after sham or SNI surgery, nalbuphine induced rate suppressant effects were less potent in both sham and SNI groups, and though there was no significant dose by surgical status interaction, this change was observed to a slightly greater extent in both male and female SNI groups compared to sham groups (Fig. 7). 4-6 months after surgery, larger doses were needed to decrease rates of responding (Fig. 7), and nalbuphine significantly decreased rates of responding in only the male sham group (Fig. 7A). Collectively, independent of surgical status, nalbuphine was less rate suppressant 4-6 months after surgery. The leftward shifts in nalbuphine-induced analgesic-like effects and rightward shifts in the rate suppressant effects were observed in both sham and SNI groups, suggesting that SNI-induced hypersensitivity did not induce changes in the potency of nalbuphine. This suggests other factors may underlie the observed shifts.

First, rats began these experiments age-matched and at approximately 8 weeks old. However, these studies are longitudinal and lasted approximately 9-11 months (see methods). Previous studies have demonstrated that nalbuphine-induced antinociceptive-like effects were more potent and efficacious in older rats (Smith & Gray, 2001). Second, it is also possible activity at other opioid receptors may contribute to the increased sensitivity to nalbuphine over time. Nalbuphine has ∼9x higher affinity for MOR than kappa opioid receptors (KOR); however, nalbuphine was more efficacious in a GTP*y*S assay at KORs (∼47%) than MORs (∼18%) (Elmariah et al., 2022). KOR agonists have been demonstrated to effectively attenuate neuropathic pain-like states in rodents (Hall et al., 2016; Paton et al., 2022). Third, nalbuphine induced rate suppressant effects decreased over time, and a decrease in behavioral disruption may explain increased sensitivity in the mechanical hypersensitivity experiments. Finally, it is somewhat unlikely that changes in nalbuphine effects are due to changes in MOR expression, as previous studies suggest a decreased expression of MORs in pain states (Back et al., 2006; Campos-Jurado et al, 2019; Dong et al., 2019; Hou et al., 2017; Ji et al., 1995; Kaneuchi et al., 2019; Pol et al., 2006; Porecca et al., 1998; Thompson et al., 2018; Yamamoto et al., 2008; Zhang et al., 1998). This would be consistent with the decreased rate suppressant effects; however, this is not consistent with increased sensitivity in Randall Selitto assays.

Previous work has demonstrated that tolerance develops differentially to various MOR agonist-induced behavioral effects, such that tolerance to analgesic and euphoric effects of MOR agonists occurs prior to tolerance to respiratory depressant or constipating actions (Angst et al., 2009; Grunkemeir 2007; Galligan et al., 2014; Mohamed et al., 2013; Paronis & Holtzman, 1994; Ross et al., 2008; Shippenberg et al., 1988). Other studies have demonstrated a different level of receptor reserve for the analgesic and primary reinforcing effects of MOR agonists (Zernig, Lewis, & Woods, 1997). Therefore, the results in the present study may suggest that nalbuphine-induced rate suppressant effects of MOR agonists have a lower receptor reserve than those of nalbuphine-induced analgesic effects.

Overall, the present study found few sex differences in MOR agonist-induced antinociceptive- and antihyperalgesic-like effects. This was somewhat unexpected as many previous studies have demonstrated sex differences (for example: Barrett, Smith, & Picker, 2002; Cook et al., 2000; Cook & Nickerson, 2005; Craft et al., 2001; Peckham & Traynor 2005, 2006; Terner et al., 2002; 2005; for review-Craft et al., 2003). However, relatively few studies have directly examined MOR agonist induced antinociceptive- and antihyperalgesic-like effects in a neuropathic model of pain and studied sex differences. Generally, more robust sex differences are observed with partial agonists and with higher intensity noxious stimuli (Cook et al., 2000).

Finally, as expected, there were group differences (sham, SNI) in paw withdrawal thresholds in Randall Selitto assays (Fig. 1B, 2); however, consistent with previous work (Wen et al., 2007), SNI induced hypersensitivity did not induce a change in latency to nocifensive behavior compared to sham animals in the hot plate assay (Fig. 3). This suggests that SNI-induced chronic nerve injury leads to mechanical hypersensitivity but not thermal hypersensitivity. It is possible that while SNI surgery produced a long-lasting mechanical hypersensitivity, the SNI animals may not be in a pain state, but rather experiencing decreases in the function of the injured limb or other sensations such as numbness in the injured hind paw/limb. This possibility is supported by lack of evidence of pain depressed operant responding in SNI groups compared to rates of responding prior to surgery and in sham groups (Fig. 6) as well as similar weight gain over time in sham and SNI groups (Fig. 1).

Though, we found that THC, diclofenac, and gabapentin produced small but significant increases in paw withdrawal thresholds in only the SNI groups (Fig. 4), which is consistent with the antihyperalgesic-like effects of these drugs. THC, diclofenac, and gabapentin did not produce significant antinociceptive-like effects in the sham groups, suggesting that these drugs did not simply disrupt behavior. Gabapentin failed to alter rates of responding in male rats; however, low doses (30 mg/kg) significantly increased rates of responding and larger doses (180 mg/kg) significantly decreased rates of responding in female rats. Pre-clinical studies are extremely useful for evaluating the effects of drugs in various experimental states; however, the present findings highlight the utility of concurrently examining behaviorally disrupting effects (e.g., rate suppression) and pain-related behaviors when using pain-elicited assays such as Randall Selitto or hot plate.

In conclusion, the present study evaluated the antinociceptive-like effects, antihyperalgesic-effects, and rate suppressant effects of opioid analgesics varying in potency and efficacy along with select non-opioid analgesics in the presence or absence of SNI-induced hypersensitivity. The overall finding was that SNI-induced chronic hypersensitivity failed to alter the antinociceptive-like effects, antihyperalgesic-like effects, and rate suppressant effects of MOR agonists. Over time, few changes were observed in the behavioral effects of higher efficacy MOR agonists, such as fentanyl or morphine. In contrast, over time, there was an increase in sensitivity to nalbuphine-induced antinociceptive-like and antihyperalgesic-like effects while a corresponding decrease in sensitivity to nalbuphine-induced rate suppressant effects was observed. Future studies should directly evaluate the abuse potential of *partial* MOR agonists in the presence or absence of chronic pain states and further characterize the antinociceptive- and anithyperalgesic-like effects of MOR agonists throughout the lifespan.

## References

Angst, M. S., Chu, L. F., Tingle, M. S., Shafer, S. L., Clark, D. J., & Drover, D. R. (2009). No evidence for the development of acute tolerance to analgesic, respiratory depressant and sedative opioid effects in humans. Pain, 142(1), 17–26.

Back, S. K., Lee, J., Hong, S. K., & Na, H. S. (2006). Loss of spinal μ-opioid receptor is associated with mechanical allodynia in a rat model of peripheral neuropathy. Pain, 123(1-2), 117–126.

Ballantyne, J. C., & LaForge, K. S. (2007). Opioid dependence and addiction during opioid treatment of chronic pain. Pain, 129(3), 235–255.

Barrett, A. C., Smith, E. S., & Picker, M. J. (2002). Sex-related differences in mechanical nociception and antinociception produced by μ-and κ-opioid receptor agonists in rats. European journal of pharmacology, 452(2), 163–173.

Bernetti, A., Agostini, F., de Sire, A., Mangone, M., Tognolo, L., Di Cesare, A., … & Paoloni, M. (2021). Neuropathic pain and rehabilitation: a systematic review of international guidelines. Diagnostics, 11(1), 74.

Bouhassira, D., Lantéri-Minet, M., Attal, N., Laurent, B., & Touboul, C. (2008). Prevalence of chronic pain with neuropathic characteristics in the general population. Pain, 136(3), 380–387. PMID: 17888574

Campos-Jurado, Y., Igual-López, M., Padilla, F., Zornoza, T., Granero, L., Polache, A., … & Hipólito, L. (2019). Activation of MORs in the VTA induces changes on cFos expression in different projecting regions: Effect of inflammatory pain. Neurochemistry International, 131, 104521.

Cavalli, E., Mammana, S., Nicoletti, F., Bramanti, P., & Mazzon, E. (2019). The neuropathic pain: An overview of the current treatment and future therapeutic approaches. International Journal of Immunopathology and Pharmacology, 33, 2058738419838383Cook & Nickerson, 2005

Cook, C. D., Barrett, A. C., Roach, E. L., Bowman, J. R., & Picker, M. J. (2000). Sex-related differences in the antinociceptive effects of opioids: importance of rat genotype, nociceptive stimulus intensity, and efficacy at the µ opioid receptor. Psychopharmacology, 150, 430–442.

Craft, R. M., Tseng, A. H., McNiel, D. M., Furness, M. S., & Rice, K. C. (2001). Receptor-selective antagonism of opioid antinociception in female versus male rats. Behavioural pharmacology, 12(8), 591–602.

Craft, R. M. (2003). Sex differences in opioid analgesia:“from mouse to man”. The Clinical journal of pain, 19(3), 175–186.

Decosterd, I., & Woolf, C. J. (2000). Spared nerve injury: an animal model of persistent peripheral neuropathic pain. Pain, 87(2), 149–158.

Dong, J., Zuo, Z., Yan, W., Liu, W., Zheng, Q., & Liu, X. (2019). Berberine ameliorates diabetic neuropathic pain in a rat model: involvement of oxidative stress, inflammation, and μ-opioid receptors. Naunyn-Schmiedeberg’s Archives of Pharmacology, 392, 1141–1149.

Dworkin, R. H., O’connor, A. B., Backonja, M., Farrar, J. T., Finnerup, N. B., Jensen, T. S., … & Wallace, M. S. (2007). Pharmacologic management of neuropathic pain: evidence-based recommendations. Pain, 132(3), 237–251.

Elmariah, S., Chisolm, S., Sciascia, T., & Kwatra, S. G. (2022). Modulation of the kappa and mu opioid axis for the treatment of chronic pruritus: a review of basic science and clinical implications. JAAD international, 7, 156–163.

Erichsen, H. K., & Blackburn-Munro, G. (2002). Pharmacological characterisation of the spared nerve injury model of neuropathic pain. Pain, 98(1-2), 151–161.

Galligan, J. J., & Akbarali, H. I. (2014). Molecular physiology of enteric opioid receptors. American journal of gastroenterology supplements (Print), 2(1), 17.

Grunkemeier, D. M., Cassara, J. E., Dalton, C. B., & Drossman, D. A. (2007). The narcotic bowel syndrome: clinical features, pathophysiology, and management. Clinical Gastroenterology and Hepatology, 5(10), 1126–1139.

Hall, S. M., Lee, Y. S., & Hruby, V. J. (2016). Dynorphin A analogs for the treatment of chronic neuropathic pain. Future Medicinal Chemistry, 8(2), 165–177.

Hayes, C. J., Krebs, E. E., Hudson, T., Brown, J., Li, C., & Martin, B. C. (2020). Impact of opioid dose escalation on pain intensity: a retrospective cohort study. Pain, 161(5), 979.

Hoffman, E. M., Watson, J. C., St Sauver, J., Staff, N. P., & Klein, C. J. (2017). Association of long-term opioid therapy with functional status, adverse outcomes, and mortality among patients with polyneuropathy. JAMA neurology, 74(7), 773–779.

Hou, Y. Y., Cai, Y. Q., & Pan, Z. Z. (2015). Persistent pain maintains morphine-seeking behavior after morphine withdrawal through reduced MeCP2 repression of GluA1 in rat central amygdala. Journal of Neuroscience, 35(8), 3689–3700.

Ji, R. R., Zhang, Q., Law, P. Y., Low, H. H., Elde, R., & Hokfelt, T. (1995). Expression of mu-, delta-, and kappa-opioid receptor-like immunoreactivities in rat dorsal root ganglia after carrageenan-induced inflammation. Journal of Neuroscience, 15(12), 8156–8166.

Joubert, F., Guerrero-Moreno, A., Fakih, D., Reboussin, E., Gaveriaux-Ruff, C., Acosta, M. C., … & Réaux-Le Goazigo, A. (2020). Topical treatment with a mu opioid receptor agonist alleviates corneal allodynia and corneal nerve sensitization in mice. Biomedicine & Pharmacotherapy, 132, 110794.

Kaneuchi, Y., Sekiguchi, M., Kameda, T., Kobayashi, Y., & Konno, S. I. (2019). Temporal and spatial changes of μ-opioid receptors in the brain, spinal cord and dorsal root ganglion in a rat lumbar disc herniation model. Spine, 44(2), 85–95.

Mohammed, W., Alhaddad, H., Marie, N., Tardy, F., Lamballais, F., Risède, P., … & Mégarbane, B. (2013). Comparison of tolerance to morphine-induced respiratory and analgesic effects in mice. Toxicology letters, 217(3), 251–259.

O’Connor, A. B., & Dworkin, R. H. (2009). Treatment of neuropathic pain: an overview of recent guidelines. The American journal of medicine, 122(10), S22–S32.

Ozaki, S., Narita, M., Narita, M., Iino, M., Sugita, J., Matsumura, Y., & Suzuki, T. (2002). Suppression of the morphine-induced rewarding effect in the rat with neuropathic pain: implication of the reduction in µ-opioid receptor functions in the ventral tegmental area. Journal of neurochemistry, 82(5), 1192–1198

Paronis, C. A., & Woods, J. H. (1997). Ventilation in morphine-maintained rhesus monkeys. II: Tolerance to the antinociceptive but not the ventilatory effects of morphine. Journal of Pharmacology and Experimental Therapeutics, 282(1), 355–362.Paton et al., 2022

Peckham, E. M., & Traynor, J. R. (2006). Comparison of the antinociceptive response to morphine and morphine-like compounds in male and female Sprague-Dawley rats. Journal of Pharmacology and Experimental Therapeutics, 316(3), 1195–1201.

Peckham, E. M., Barkley, L. M., Divin, M. F., Cicero, T. J., & Traynor, J. R. (2005). Comparison of the antinociceptive effect of acute morphine in female and male Sprague–Dawley rats using the long-lasting mu-antagonist methocinnamox. Brain research, 1058(1-2), 137–147.

Pol, O., Murtra, P., Caracuel, L., Valverde, O., Puig, M. M., & Maldonado, R. (2006). Expression of opioid receptors and c-fos in CB1 knockout mice exposed to neuropathic pain. Neuropharmacology, 50(1), 123–132.

Raja, S. N., Carr, D. B., Cohen, M., Finnerup, N. B., Flor, H., Gibson, S., … & Vader, K. (2020). The revised International Association for the Study of Pain definition of pain: concepts, challenges, and compromises. Pain, 161(9), 1976–1982.

Rice, A. S., Smith, B. H., & Blyth, F. M. (2016). Pain and the global burden of disease. Pain, 157(4), 791–796.

Ross, G. R., Gabra, B. H., Dewey, W. L., & Akbarali, H. I. (2008). Morphine tolerance in the mouse ileum and colon. Journal of Pharmacology and Experimental Therapeutics, 327(2), 561–572.

Shippenberg, T. S., Emmett-Oglesby, M. W., Ayesta, F. J., & Herz, A. (1988). Tolerance and selective cross-tolerance to the motivational effects of opioids. Psychopharmacology, 96, 110–115.

Smith, M. A., & Gray, J. D. (2001). Age-related differences in sensitivity to the antinociceptive effects of opioids in male rats: influence of nociceptive intensity and intrinsic efficacy at the mu receptor. Psychopharmacology, 156, 445–453.Fentanyl morphine hot plate study

Terner, J. M., Barrett, A. C., Grossell, E., & Picker, M. J. (2002). Influence of gonadectomy on the antinociceptive effects of opioids in male and female rats. Psychopharmacology, 163(2), 183–193

Terner, J. M., Lomas, L. M., & Picker, M. J. (2005). Influence of estrous cycle and gonadal hormone depletion on nociception and opioid antinociception in female rats of four strains. The journal of pain, 6(6), 372–383.

Thompson, S. J., Pitcher, M. H., Stone, L. S., Tarum, F., Niu, G., Chen, X., … & Bushnell, M. C. (2018). Chronic neuropathic pain reduces opioid receptor availability with associated anhedonia in rat. Pain, 159(9), 1856.

Van Hecke, O. A. S. K. R., Austin, S. K., Khan, R. A., Smith, B. H., & Torrance, N. (2014). Neuropathic pain in the general population: a systematic review of epidemiological studies. PAIN®, 155(4), 654–662.

Walker, E. A., & Young, A. M. (1993). Discriminative-stimulus effects of the low efficacy mu agonist nalbuphine. The Journal of pharmacology and experimental therapeutics, 267(1), 322–330.

Widerström-Noga, E. G., & Turk, D. C. (2003). Types and effectiveness of treatments used by people with chronic pain associated with spinal cord injuries: influence of pain and psychosocial characteristics. Spinal cord, 41(11), 600–609

Yamamoto, J., Kawamata, T., Niiyama, Y., Omote, K., & Namiki, A. (2008). Down-regulation of mu opioid receptor expression within distinct subpopulations of dorsal root ganglion neurons in a murine model of bone cancer pain. Neuroscience, 151(3), 843–853.

Zernig, G., Lewis, J. W., & 229. Woods, J. H. (1997). Clocinnamox inhibits the intravenous self-administration of opioid agonists in rhesus monkeys: comparison with effects on opioid agonist-mediated antinociception. Psychopharmacology, 129, 233–242.

Zhang, Q., Schäfer, M., Elde, R., & Stein, C. (1998). Effects of neurotoxins and hindpaw inflammation on opioid receptor immunoreactivities in dorsal root ganglia. Neuroscience, 85(1), 281–291.

